# Fast and scalable querying of eukaryotic linear motifs with *gget elm*

**DOI:** 10.1101/2023.11.15.567056

**Authors:** Laura Luebbert, Chi Hoang, Manjeet Kumar, Lior Pachter

## Abstract

**Motivation:** Eukaryotic linear motifs (ELMs), or Short Linear Motifs (SLiMs), are protein interaction modules that play an essential role in cellular processes and signaling networks and are often involved in diseases like cancer. The ELM database is a collection of manually curated motif knowledge from scientific papers. It has become a crucial resource for cataloging motif biology and recognizing candidate ELMs in novel amino acid sequences. Users can search amino acid sequences or UniProt IDs on the ELM resource web interface. However, as with many web services, there are limitations in the swift processing of large-scale queries through the ELM web interface or API calls, and, therefore, integration into protein function analysis pipelines is limited.

**Results:** To allow swift, large-scale motif analyses on protein sequences using ELMs curated on the ELM database, we have developed a Python and command line tool, *gget elm*, which relies on local computations for efficiently finding candidate ELMs in user-submitted amino acid sequences and UniProt identifiers. *gget elm* increases accessibility to the information stored in the ELM database and allows scalable searches for motif-mediated interaction sites in the amino acid sequences.

**Availability and implementation:** The manual and source code are available at https://github.com/pachterlab/gget.

## Introduction

Eukaryotic linear motifs (ELMs), also known as Short Linear Motifs (SLiMs), are short stretches of contiguous amino acids, typically 3 to 15 residues in length, encoding protein-protein interaction sites. They are mainly located in the intrinsically disordered regions (IDRs) of proteins and are typically found to be highly conserved in orthologous proteins. These modules can encode multiple functionalities, which include modification, degradation, docking, targeting, and binding sites for protein domains. As such, ELM-mediated interactions play an essential role in cellular processes and signaling networks, including the regulation of homeostasis, apoptosis, and differentiation (Van Roey *et al*., 2014; Davey *et al*., 2012). Pathogens like SARS-CoV2 mimic ELMs to gain entry into the cell (Kruse *et al*., 2021; Mészáros *et al*., 2021), and mutations in sequences containing ELMs contribute to diseases like cancer (Uyar *et al*., 2014; Mészáros *et al*., 2017). As a result, ELM-mediated protein interactions are potential targets for therapeutic intervention (Mészáros *et al*., 2021; Simonetti *et al*., 2023; Fasano *et al*., 2022).

The ELM resource has two main components: an exploratory candidate motif search web interface and a database with manually curated linear motif knowledge, including information on binding partners and recognition features along with associated biological context. The database information is derived from the scientific literature by expert ELM curators who analyze motif-containing sequences to capture key insights, such as the residues involved in the interaction, their evolutionary conservation, local sequence context in flanking regions, features of the binding site on the interacting partner, and other motif-related insights. In addition, the curation process captures relevant information on the contextual knowledge, which includes cellular function, location, and taxonomic distribution of motif-containing proteins. Since the database was first created (Puntervoll *et al*., 2003; Dinkel *et al*., 2011), it has been continuously updated and has been widely used for both biomedical studies as well as basic research molecular studies (Kumar *et al*., 2020, 2022; Gouw *et al*., 2018; Dinkel *et al*., 2015; Carberry, 2008; Kumar *et al*., 2023). Users can search amino acid sequences or UniProt IDs on the ELM database web interface (http://elm.eu.org/) or by submitting an API request through the ELM server. However, these methods have processing limitations when performing large-scale queries, and many requests being submitted simultaneously can lead to server overload and extended wait times.

To expedite ELM investigation, we have developed a Python and command line tool relying on local computations for efficiently finding ELMs in user-submitted amino acid sequences or UniProt IDs: *gget elm*. Our tool extends the *gget* suite of tools (Luebbert and Pachter, 2022). *gget elm* increases accessibility to the information stored in the ELM database and allows scalable searches for ELMs in amino acid sequences. The command line interface and optional JSON formatted output allow swift integration into existing protein analysis workflows.

## Description

Users can submit an amino acid sequence or a UniProt ID to *gget elm. gget elm* captures both homology-based matches corresponding to curated motifs in orthologous proteins in the ELM database and POSIX regular expression (regex) matches corresponding to candidate motifs in the provided sequence. Hence, *gget elm* returns two separate data frames (or JSON formatted dictionaries for use from the command line) containing the respective motif matches and extensive information about each motif. Figure 1 provides an overview of the *gget elm* back-end.

**Figure 1.**
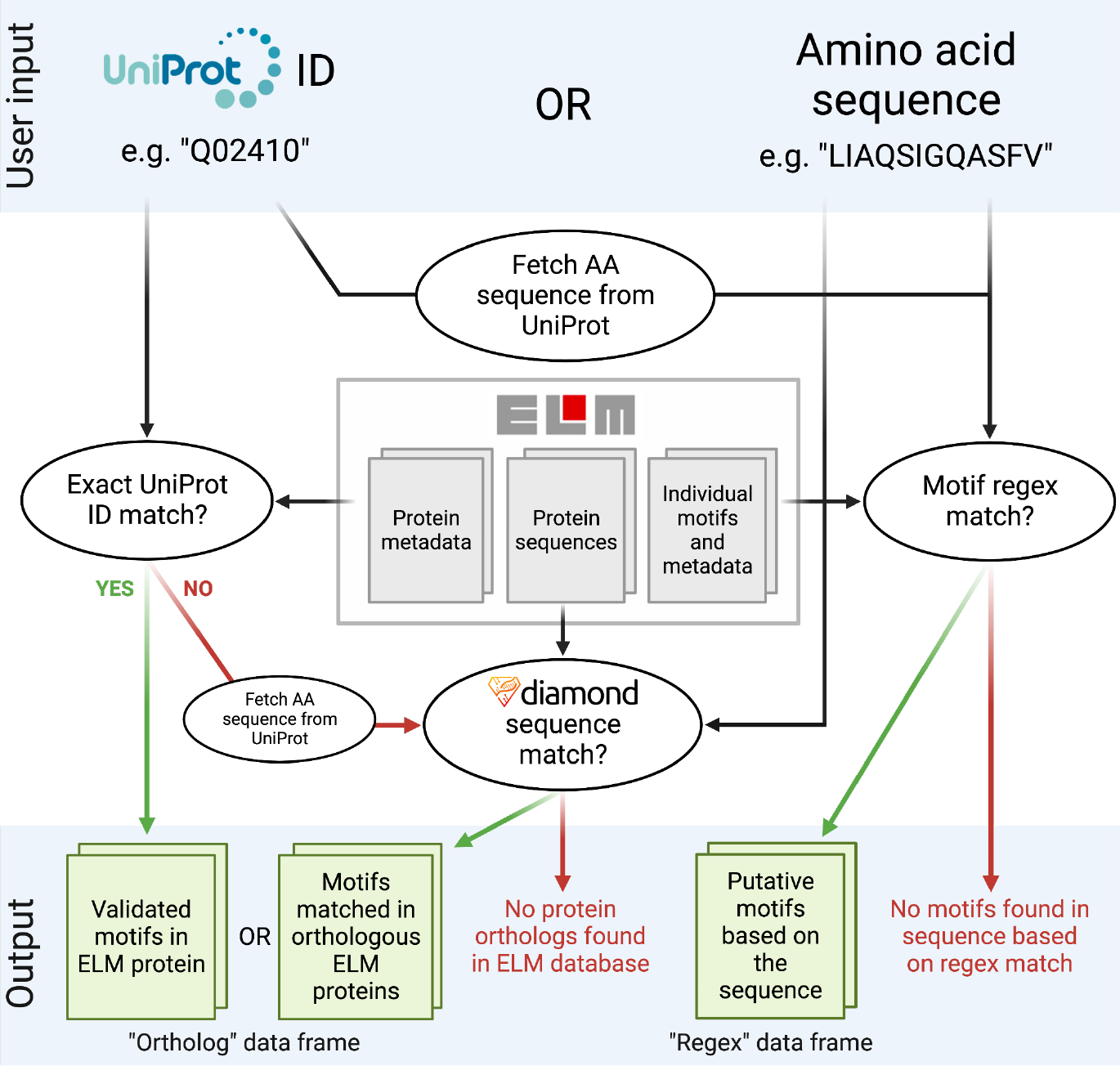
Schematic overview of the *gget elm* back-end.

After installing *gget* ($ pip install gget), the user downloads the ELM database reference information using a specialized module, *gget setup*, with the command $ gget setup elm. This command may be repeated at any time to update the local copy of the ELM database, which currently requires a total of 3 MB of disk space. The files are saved in the *gget* installation directory.

If the user submits a UniProt ID to *gget elm* and the protein is not present in the ELM database, its amino acid sequence is fetched from UniProt (UniProt Consortium, 2021). Using the DIAMOND alignment algorithm (Buchfink *et al*., 2021), the sequence is compared to the motif-containing proteins in the ELM database. *gget elm* returns all motifs associated with orthologous proteins, including information about each orthologous protein, and extensive details on each motif. *gget elm* also returns alignment scores for each DIAMOND hit, including identity and coverage percentages and boolean output on whether the orthologous motif is contained within the overlapping region between the query and subject sequence. To compute the regex data frame, *gget elm* considers all regex expressions from the ELM database and scans them against the provided amino acid sequence to report all matches. The data from the ELM database is combined to return relevant information about each matched interaction motif, including motif description, type, sequence, location in the ortholog and query sequence, and host taxonomy, for both data frames.

*gget elm* builds on existing *gget* modules, such as *gget seq* to fetch amino acid sequences from UniProt, and a new module developed in parallel with *gget elm*: *gget diamond*, which aligns sequences using the DIAMOND algorithm (Buchfink *et al*., 2021) and can be used independently from *gget elm*.

While *gget elm* results are similar to results obtained through the ELM web interface, they may not be identical due to differences underlying the computations. For example, *gget elm* uses DIAMOND for fast and sensitive local alignment of the amino acid sequences, whereas the ELM web interface has its own suite of back-end tools and deliberately limits the number of proteins in the output to be manageable for the web server (Chica *et al*., 2008). Some exemplary results obtained by *gget elm* are explored in this Google Colab notebook: https://tinyurl.com/4bd5h8hr.

### Usage and Documentation

Akin to all modules contained within *gget* (Luebbert and Pachter, 2022), *gget elm* features an extensive manual available as function documentation in a Python environment or as standard output using the help flag [-h] in the command line. The accuracy of the returned results is maintained through extensive unit tests, which automatically run on a bi-weekly basis. The complete manual with examples can be viewed on the *gget* website in English (https://pachterlab.github.io/gget/en/elm) and in Spanish (https://pachterlab.github.io/gget/es/elm).

*gget* can be installed from PyPI using the command line with the following command:

~~~
$ pip install gget
~~~

Alternatively, *gget* can be installed using Anaconda:

~~~
$ conda install -c bioconda gget
~~~

Example *gget elm* commands to find ELMs in a protein from its amino acid sequence or UniProt ID look as follows:

Command line (JSON formatted results are saved in a folder named ‘results’):

~~~
$ gget setup elm # Downloads/updates local ELM database
$ gget elm -o results LIAQSIGQASFV
$ gget elm -o results --uniprot Q02410
~~~

Python (two data frames are returned):

~~~
>>> gget.setup(“elm”) # Downloads/updates local ELM database
>>> ortholog_df, regex_df = gget.elm(“LIAQSIGQASFV”)
>>> ortholog_df, regex_df = gget.elm(“Q02410”, uniprot=True)
~~~

The [--threads] argument can be used to multithread the sequence alignment for increased speed for large-scale computations.

### Proof of concept: *gget elm* reports the loss of a protein interaction motif involved in DNA repair in a carcinogenic BRCA2 mutation

BRCA2 (BReast CAncer gene 2) plays an essential role in DNA repair through homologous recombination, and heterozygous germline defects in BRCA2 increase the risk of breast cancer. The promotion of homologous recombination by BRCA2 requires its association with the partner and localizer of BRCA2 (PALB2) (Hanenberg and Andreassen, 2018). This important protein-protein interaction occurs at the site of a linear motif (ELM: LIG_PALB2_WD40_1, regex: [WF..L]), which can be recognized by *gget elm*. In the experiment shown in Supplementary Figure 1, we analyze the wildtype BRCA2 sequence and a mutant BRCA2 sequence with a single amino acid substitution (W31C), previously described as carcinogenic due to a loss of interaction with PALB2 (Oliver *et al*., 2009). *gget elm* accurately reports the loss of the PALB2 interaction motif in the mutant sequence compared to the wildtype sequence (Supplementary Figure 1, https://tinyurl.com/yc5r2b5m).

## Discussion

We have shown that *gget elm* facilitates scalable querying of the ELM database via local queries, and its use via the command line makes it easy to integrate into scripted workflows. While this feature should extend the usability of the ELM database, there are limitations while performing motif searches using the ELM database web interface or *gget elm*. A common problem encountered is that short and degenerate ELMs inevitably lead to false positive matches. Accuracy can be improved by filtering the results using the additional contextual information, which is also returned by *gget elm*, including description, structural features, and host taxonomy. Furthermore, combining motif results with structural and alignment information can provide information about the functional availability of the interaction site (Lee *et al*., 2023). The 3D structure of a protein can be predicted from its amino acid sequence *de novo* using algorithms like AlphaFold2 (Jumper *et al*., 2021) and compared to experimentally derived crystal structures of orthologs deposited on the PDB (Berman *et al*., 2000). The *gget* suite of tools contains a workflow to perform both of these computations, which is demonstrated here: https://tinyurl.com/yzc9ytvx.

## Author contributions

LL and LP conceived the project after listening to a lecture by Prof. Amy E. Keating. LL, CH, and MK designed the *gget elm* approach. LL and CH wrote the *gget elm* software, with CH being the primary developer under the supervision of LL. LL is the primary developer of the *gget* software, and MK is the primary developer of the ELM resource. LL wrote the initial draft of the manuscript. CH, MK, and LP provided feedback on the manuscript. All authors reviewed and approved the manuscript.

## Acknowledgments

We thank the expert curators of the ELM database for providing an excellent resource. We thank Dr. Toby Gibson for the valuable feedback on the manuscript. We also thank Candace Rypisi and the rest of the Summer Undergraduate Research Fellowships (SURF) program staff for facilitating valuable research opportunities for undergraduate students and mentorship opportunities for graduate students at Caltech. Illustrations in Figure 1 were created with BioRender.com.

## Funding

LL was supported by funding from the Biology and Bioengineering Division at the California Institute of Technology and the Chen Graduate Innovator Grant CHEN.SYS3.CGIAFY21. CH was supported by the Citadel Global Fixed Income SURF Fellowship. *gget* was supported by Pachter lab start-up funds. *Conflict of Interest:* none declared.

## Notes

https://pachterlab.github.io/gget/

### Competing Interest Statement

The authors have declared no competing interest.

