## Supplementary Figure 1 for "Fast and scalable querying of eukaryotic linear motifs with *gget elm*"

### `gget elm` predicts the loss of a protein interaction motif involved in DNA repair in a carcinogenic BRCA2 mutation

Reference: Oliver AW, Swift S, Lord CJ, Ashworth A, Pearl LH. Structural basis for recruitment of BRCA2 by PALB2. EMBO Rep. 2009 Sep;10(9):990-6. doi: 10.1038/embor.2009.126. Epub 2009 Jul 17. Erratum in: EMBO Rep. 2017 Jul;18(7):1264. PMID: 19609323; PMCID: PMC2750052. https://www.ncbi.nlm.nih.gov/pmc/articles/PMC2750052/

ELM data can be downloaded & distributed for non-commercial use according to the ELM Software License Agreement.

Install `gget` and set up `gget elm`:

```
!pip install -qU gget
```

```
     ━━━━━━━━━━━━━━━━━━━━━━━━━━━━━━━━━━━━━━━━ 25.2/25.2 MB 21.1 MB/s eta 0:00:00
  Preparing metadata (setup.py) ... done
  Building wheel for gget (setup.py) ... done
```

```
import gget
gget.setup("elm")
```

```
      % Total    % Received % Xferd  Average Speed   Time    Time     Time  Current
                                     Dload  Upload   Total   Spent    Left  Speed
   %   0     0    0     0    0     0      0      0 --:--:-- --:--:-- --:--:--     0Total    % Received % Xferd  Average Speed
                                     Dload  Upload   Total   Spent    Left  Speed
   100 87336  100 87336    0     0  91237      0 --:--:-- --:--:-- --:--:-- 91164
     % Total    % Received % Xferd  Average Speed   Time    Time     Time  Current
                                     Dload  Upload   Total   Spent    Left  Speed
   100  914k  100  914k    0     0   405k      0  0:00:02  0:00:02 --:--:--  405k
   100 2123k  100 2123k    0     0   635k      0  0:00:03  0:00:03 --:--:--  635k
```

Find protein interaction motifs based on ortholog and regex matches in the wildtype and mutant BRCA2 amino acid sequences:

```
# Define wildtype and mutant BRCA2 amino acid sequences
wt_brca2 = "MPIGSKERPTFFEIFKTRCNKADLGPISLNWFEELSSEAPPYNSEPAEESEHKNNNYEPNLFKTPQRKPSYNQLASTPIIFKEQGLTLPLYQSPVKELDKFKLDLGRNVPNSRHKSL
mutant_brca2 = "MPIGSKERPTFFEIFKTRCNKADLGPISLNCFEELSSEAPPYNSEPAEESEHKNNNYEPNLFKTPQRKPSYNQLASTPIIFKEQGLTLPLYQSPVKELDKFKLDLGRNVPNSR

# Visualize the single amino acid difference between the sequences (W31C)
gget.muscle([wt_brca2, mutant_brca2])
```

```
Seq0
        MPIGSKERPTFFEIFKTRCNKADLGPISLNWFEELSSEAPPYNSEPAEESEHKNNNYEPNLFKTPQRKPSYNQLASTPII
Seq1
        MPIGSKERPTFFEIFKTRCNKADLGPISLNCFEELSSEAPPYNSEPAEESEHKNNNYEPNLFKTPQRKPSYNQLASTPII

Seq0
        FKEQGLTLPLYQSPVKELDKFKLDLGRNVPNSRHKSLRTVKTKMDQADDVSCPLLNSCLSESPVVLQCTHVTPQRDKSVV
Seq1
        FKEQGLTLPLYQSPVKELDKFKLDLGRNVPNSRHKSLRTVKTKMDQADDVSCPLLNSCLSESPVVLQCTHVTPQRDKSVV

Seq0
        CGSLFHTPKFVKGRQTPKHISESLGAEVDPDMSWSSSLATPPTLSSTVLIVRNEEASETVFPHDTTANVKSYFSNHDESL
Seq1
        CGSLFHTPKFVKGRQTPKHISESLGAEVDPDMSWSSSLATPPTLSSTVLIVRNEEASETVFPHDTTANVKSYFSNHDESL

Seq0
        KKNDRFIASVTDSENTNQREAASHGFGKTSGNSFKVNSCKDHIGKSMPNVLEDEVYETVVDTSEEDSFSLCFSKCRTKNL
Seq1
        KKNDRFIASVTDSENTNQREAASHGFGKTSGNSFKVNSCKDHIGKSMPNVLEDEVYETVVDTSEEDSFSLCFSKCRTKNL

Seq0
        QKVRTSKTRKKIFHEANADECEKSKNQVKEKYSFVSEVEPNDTDPLDSNVANQKPFESGSDKISKEVVPSLACEWSQLTL
Seq1
        QKVRTSKTRKKIFHEANADECEKSKNQVKEKYSFVSEVEPNDTDPLDSNVANQKPFESGSDKISKEVVPSLACEWSQLTL
```

```
Seq0
        SGLNGAQMEKIPLLHISSCDQNISEKDLLDTENKRKKDFLTSENSLPRISSLPKSEKPLNEETVVNKRDEEQHLESHTDC
Seq1
        SGLNGAQMEKIPLLHISSCDQNISEKDLLDTENKRKKDFLTSENSLPRISSLPKSEKPLNEETVVNKRDEEQHLESHTDC

Seq0
        ILAVKQAISGTSPVASSFQGIKKSIFRIRESPKETFNASFSGHMTDPNFKKETEASESGLEIHTVCSQKEDSLCPNLIDN
Seq1
        ILAVKQAISGTSPVASSFQGIKKSIFRIRESPKETFNASFSGHMTDPNFKKETEASESGLEIHTVCSQKEDSLCPNLIDN

Seq0
        GSWPATTTQNSVALKNAGLISTLKKKTNKFIYAIHDETSYKGKKIPKDQKSELINCSAQFEANAFEAPLTFANADSGLLH
Seq1
        GSWPATTTQNSVALKNAGLISTLKKKTNKFIYAIHDETSYKGKKIPKDQKSELINCSAQFEANAFEAPLTFANADSGLLH

Seq0
        SSVKRSCSQNDSEEPTLSLTSSFGTILRKCSRNETCSNNTVISQDLDYKEAKCNKEKLQLFITPEADSLSCLQEGQCEND
Seq1
        SSVKRSCSQNDSEEPTLSLTSSFGTILRKCSRNETCSNNTVISQDLDYKEAKCNKEKLQLFITPEADSLSCLQEGQCEND

Seq0
        [illegible]
```

```
# Predict protein interaction domains in wildtype sequence
wt_ortho, wt_regex = gget.elm(wt_brca2)
```

```
# Predict protein interaction domains in mutant sequence
mutant_ortho, mutant_regex = gget.elm(mutant_brca2)
```

The BRCA2 interaction motif required for the interaction with PALB2 which plays a role in DNA repair is lost in the carcinogenic mutant:

```
# Do not truncate data frames
import pandas as pd
pd.set_option('display.max_colwidth', None)
```

```
# Show wildtype interactions domains involved in DNA repair
wt_regex[wt_regex["Description"].str.contains("DNA double strand breaks and repair")]
```

| Instance_accession | ELMIdentifier | FunctionalSiteName | ELMType | Descri |
|---|---|---|---|---|

```
# These motifs are lost in the mutant sequence
mutant_regex[mutant_regex["Description"].str.contains("DNA double strand breaks and repair")]
```
